# Where and why alongshore variation in larval transport enables the establishment of introduced species

**DOI:** 10.64898/2026.08.14.744914

**Authors:** James M. Pringle, William G. Lush, James E. Byers

## Abstract

After introduction, many non-native marine species are dispersed planktonically. Secondary spread within the non-native range has been shown to prevent the establishment of the introduced species if the advection of larvae prevents sufficient return of larvae to maintain the population in the face of competition with native species. However, those studies have largely neglected the effects of spatial variation in alongshore larval transport.

We examine the introduction of a novel species with planktonic dispersal into a more realistic coastal environment which includes spatial variation in larval transport estimated from the Mercator Ocean 1/12^th^ degree global circulation model. The introduction may either be from a distant habitat, or through range expansion.

We find that there are locations in the global coastal ocean where introduced species are more likely to persist because of spatial variation of coastal currents. These include regions where alongshore larval transport diverges, such as estuaries. The location where a non-native species is introduced may not be where it flourishes – it cannot be assumed that the region where invading species are first noticed to be abundant is the region where it was introduced. We extend closed-population theory to open coastal systems to estimate persistence as a function of local circulation, habitat extent, and the competitive advantage of the introduced species.

Software is provided which allows the estimations of regions where introduced species are more likely to persist and flourish as a function of larval depth behavior, planktonic duration and release timing.

## Introduction

Understanding where non-native species can be established along the coastal ocean and what controls their success is vital to understanding and managing coastal environments (Mooney and Drake 2012; Byers et al. 2015). For a non-native species to flourish when introduced into a new region, it must persist in a region after the initial transport of colonizers to a new region. Many introductions fail at this point (Carlton 1996). Understanding what determines the success of these invasion attempts has been the focus of efforts over the past decades (Roy et al. 2002; Johnston et al. 2009; Zhan et al. 2015; Papacostas et al. 2017; Geburzi and McCarthy 2018). For many introduced species, including benthic invertebrates, sea-grasses and seaweeds, a principal mechanism for post-introduction dispersal (also known as secondary spread) is planktonic dispersal of some sort – either through larvae, rafting, or other similar dispersal mechanisms mediated by ocean currents (Strathmann 1985; Muhlin et al. 2008; Johnson et al. 2012).

In coastal oceans there are alongshore flows which will transport larvae or other propagules on average in a predictable direction, although with considerable variation around the mean direction of transport (Brink 2005; Lush and Pringle 2025). The mean transport of propagules away from the location of the parents makes the successful persistence of the invader where introduced more difficult because larvae are lost offshore or to downstream transport (Byers and Pringle 2006). Byers and Pringle (2024) find that an introduced species must have a large realized reproductive success after competition with native species in order to successfully persist and invade. However, their results are based on the assumption that the ability of newly introduced species to persist is not influenced by any along-shore variation in ocean currents.

Regional studies (Pappalardo et al. 2015; Pringle et al. 2017) and idealized modeling (Pringle and Wares 2007) suggest, however, that alongshore variation in the ocean currents or alongshore variation in habitat can make it easier for a newly introduced potential invader to persist at some locations along a coast, while making it more difficult for them to persist at others.

We develop qualitative and quantitative explanations for why alongshore variation in larval transport can alter the vulnerability of different regions to new introduced species. This extends prior work by including both alongshore variation in larval transport (e.g. Byers and Pringle 2024) and oceanographically realistic variability in ocean currents (e.g. Gaylord and Gaines 2000). To focus on the effects of dispersal, this work assumes that all competing species have the same dispersal and phenology, their competition is solely through their neutral competition for habitat, and their relative competitiveness is only expressed through different reproductive output. These assumptions lead to the simplest system that can capture the effects of alongshore variation in larval transport.

In the methods, an idealized population model which models the distribution of multiple species with potentially different reproductive output but similar dispersal strategies is described. In the results, the model is used to: 1) Describe the spatial patterns of alongshore variation of alongshore larval transport in an idealized linear coastal habitat which make a given region of coastline more susceptible (“invasible”) to the introduction of novel species. 2) Qualitatively describe how the results of 1 can be used to explain which parts of an actual coast (in this case the coast of the Americas) are more invasible for a given combination of larval duration, behavior, and phenology. 3) Quantify the likelihood that a given level of propagule pressure will lead to a successful introduction as a function of the local oceanographic conditions, population density, and relative competitive advantage of the introduced species.

These results are then used to describe the importance of the relative competitive advantage of introduced non-native species to their persistence, why estuaries are often more vulnerable to invasions, and how locations which are susceptible to invasions are also important stepping-stones for species shifting their range in response to climate change. We then describe the limitations of the results caused by the limited resolution of extant ocean models and the simplicity of our population model. Software is provided in the online supplement to reproduce these results for different regions, planktonic phenology and vertical behavior (https://github.com/JamiePringle/PringleLushByers_InvasibilityInRealOcean), and we illustrate how the software can be configured using an example that describes the northward invasion of *Carcinus maenas* along the east coast of North America.

## Methods: The modeling framework

All model results presented below are based on a simple population model. In this model, there are a finite number of model “patches” whose population is limited to *P_max_* individuals. Adults in these patches can be of an arbitrary number of different species. For this work, each species has the same dispersal ability, but can have a varying fecundity. Species reproduce asexually, so each offspring has a single unique parent, and it is of the same species (i.e. there is no speciation). Each adult produces a fixed number of dispersing propagules, hereafter referred to as larvae. The species are semelparous with non-overlapping generations, and all reproduce at the same time. The species’ larvae are equally competitive for habitat, so if more than *P_max_* larvae arrive at a patch, the successful recruits are chosen randomly. This is a model of density dependent limitation, and the only difference in the relative competitiveness of these species is in their reproductive output. There is no other competition for resources (e.g., food is assumed not be limiting, and there is no predation of adults). The model will be run in two forms; Idealized model runs on a linear coast will illustrate the basic dynamics of introduction and persistence of species, while model runs parameterized by realistic numerical models of ocean circulation will illustrate how the results apply to actual coastlines.

### Idealized model

One set of model runs is done in an idealized one-dimensional domain which is 1024 patches long and 1 patch wide. Each patch is 1km in size, and *P_max_* is 1, so that the total population across the full domain is roughly 1000 individuals (stochastically, some patches may receive no larvae and be un-occupied for a generation). As in Byers & Pringle (2006), the larvae are assumed to move an average distance L_adv_ downstream of their parents, but the dispersal distance for each individual varies around this mean with a standard deviation of L_diff_ and a Gaussian distribution. Each adult produces 10 larvae. If the larvae disperse within the domain, they can compete for habitat. If they are dispersed beyond the domain, they die. For all choices of L_adv_ and L_diff_ presented below, the larval production rate is sufficient to maintain the population in the domain (Byers and Pringle 2006). In all runs, L_diff_ is 50km. Runs are made with L_adv_=0, L_adv_=50km uniformly in one direction (left to right in Figure **1**), L_adv_=50km but converging towards the center of the domain (i.e., towards the right on the left side of the domain, and towards the left on the right side of the domain), and L_adv_=50km but diverging from the center of the domain (to the right on the right side of the domain, and to the left on the left side of the domain). These dispersal patterns and the model runs are shown in Figure **1**.

**Figure 1:**
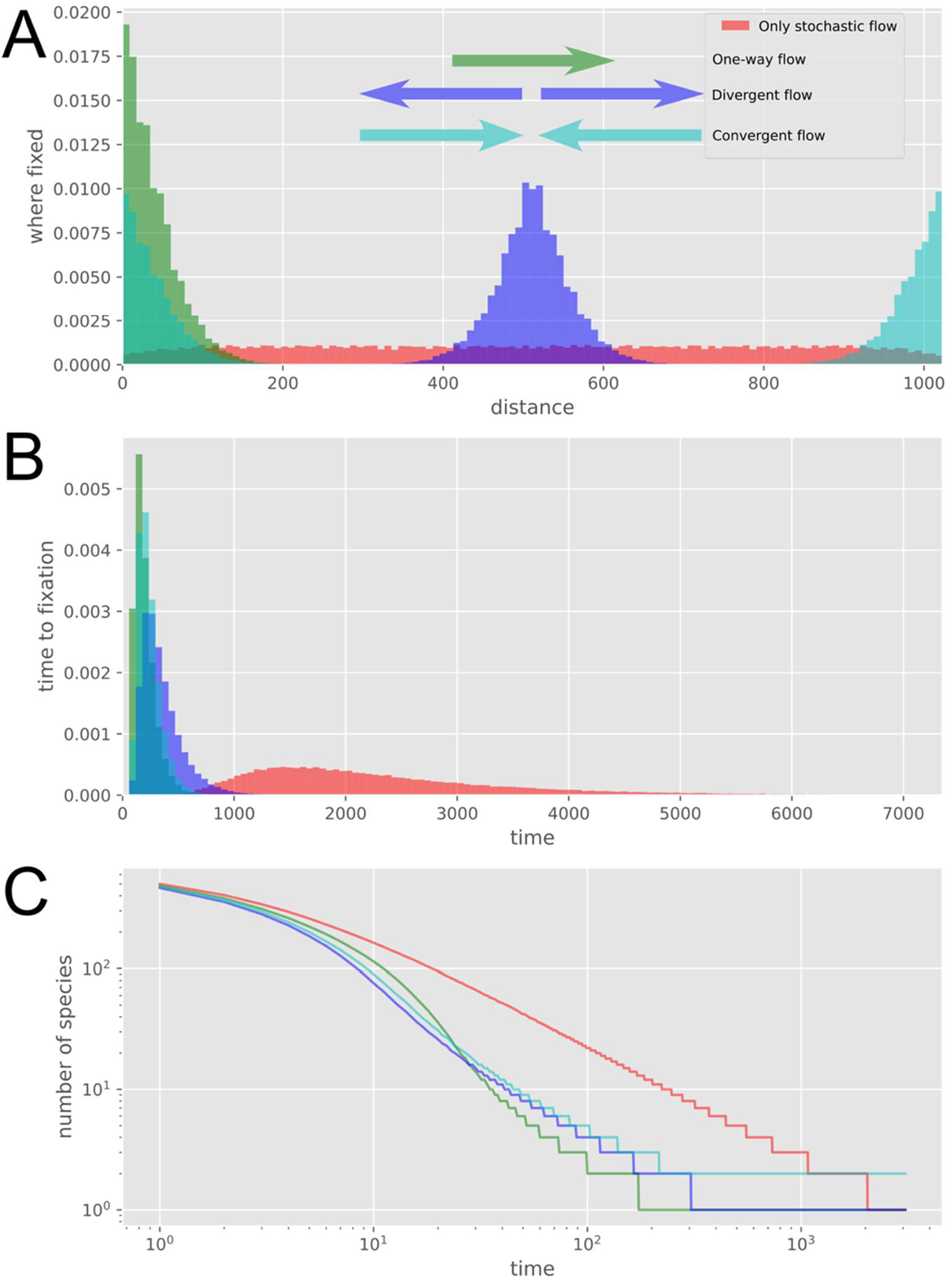
A) The starting location of the species that eventually grows to fixation by becoming the only species in the habitat (or, in the case of the converging flow, the last two species in the habitat). The colored arrows indicate the nature of the mean larval transport. The stochastic component of larval transport is the same in all cases. B) Histograms of the time until one species reaches fixation (or, in the case of converging flow, two species). C) The mean number of species as a function of time for each flow condition.

### Models with more realistic dispersal, geometry, and different species competitiveness

Another set of model runs was made with a habitat geometry and connectivity that approximates the nearshore of North and South America. The larval dispersal and habitat distribution is based on the 1/12^th^ of a degree Mercator GLORYS V12.1. (1/12^th^ of a degree is 9km at the equator, and 5km at 60N). GLORYS is a data assimilative ocean circulation model forced with surface fluxes from numerical atmospheric models and assimilates multiple sources of in situ and remotely sensed ocean data to get an optimal estimate of ocean currents (Cummings and Smedstad 2013; Lellouche et al. 2018). It includes wind- and river-driven current variability, and resolves inter-annual variability on El Nino scales (Amaya et al. 2023). This model has been shown to outperform or match other global models in predicting coastal currents (Wilkin and Hunter 2013; Amaya et al. 2023). Lush and Pringle (2025) compared Lagrangian paths calculated in the global coastal ocean with GLORYS surface currents to those observed by drifters from the Ocean Drifter Project. They found that the mean Lagrangian path was very well predicted, but that there was moderate under-prediction of the stochastic component of transport. Precomputed Lagrangian pathways are taken from the EZfate package developed by J. Pringle (https://github.com/JamiePringle/EZfate), and were computed by the Ocean Parcels software package v.2.2 (Delandmeter and Van Sebille 2019). The main limitations of this modeling product are this reduced variability and the relatively coarse resolution compared to the size of important estuaries (in Figure **2**C the model gridpoints are shown as dots in and around the Chesapeake estuary). The implications of these limitations are discussed below.

**Figure 2:**
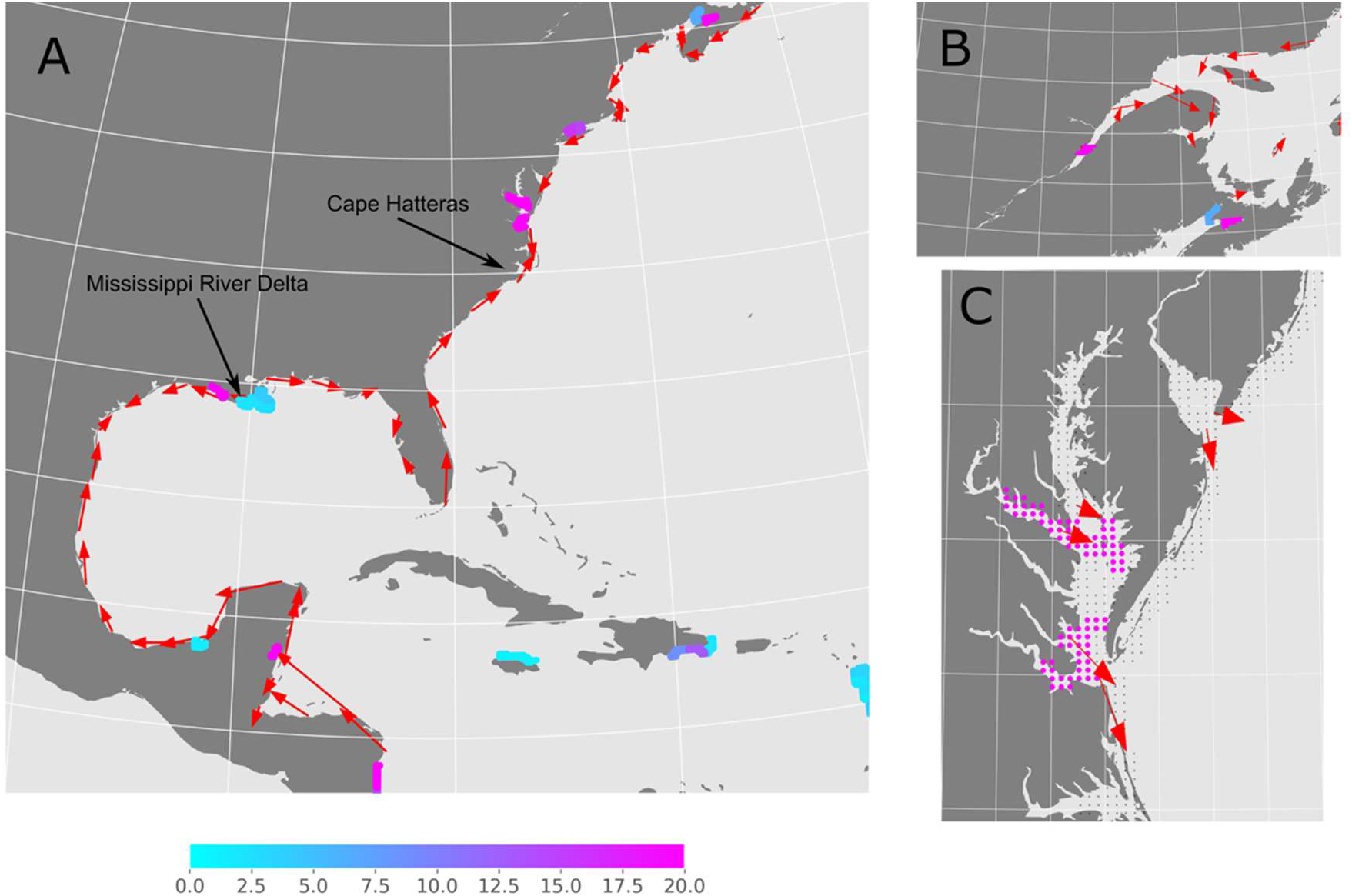
A) For larvae dispersing for 22 days at the ocean surface and released from April through June of 2007 to 2023, the direction and distance of mean larval transport at selected points along the coast of North and Central America (arrows), and the fraction of introductions (in percent) which persist for 600 generations somewhere in the domain, plotted as a colored dot at location of introduction. For this result, 10 neutral individuals are introduced into a domain where each patch has a carrying capacity of 1 individual. Statistics computed over 1000 runs. B) same as (A) but focused on the Laurentian Estuary and with selected arrows to show estuarine circulation. C) same as (B) but for Chesapeake and Delaware estuaries. The grey dots in C represent the distribution of the habitat grid, which is the same as the density grid for the Mercator GLORYS 1/12^th^ of a degree model.

The habitat of this model is meant to approximate that of nearshore coastal organisms, and so patches are defined as the model gridpoints (centered on the density grid) within 2 gridpoints (1/6^th^ of a degree) of land. The habitat is limited to North and South America south of 60°N. Patches are shown as dots in Figure **3**B, and the entire habitat is shown in figure **3**A. The carrying capacity of the habitat patches, *P_max_*, is varied in different model runs between 1 and 10 individuals per patch. The carrying capacity of 1 to 10 individuals per 1/12^th^ degree square region is kept relatively low to keep the model computationally tractable. The extension of the model results to greater population densities is discussed below.

**Figure 3:**
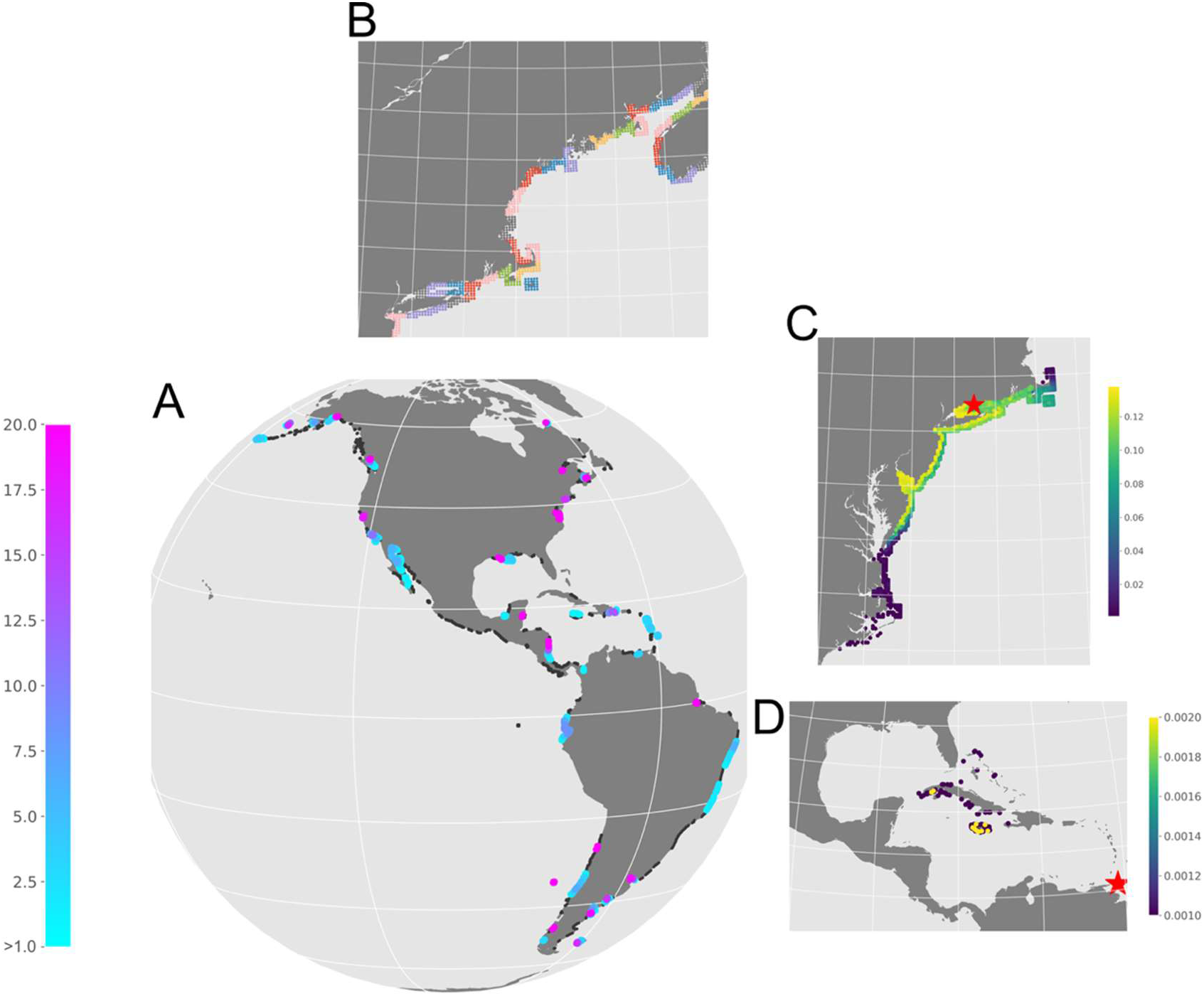
A) The likelihood of persistence for 600 generations of a neutral introduced species for locations along the coasts of North and South America. The larvae drift at the surface and are in the plankton for 22 days and connectivity is calculated for releases from April through June of 2007 to 2023. Fraction calculated for 1000 trial introductions, patch carrying capacity *P_max_*=1, *n_intro_=*10 introductions per trial. Dots are colored if the percentage which persist is >1%, and are grey if between >0% and <1%. B) The different color patches show the size of the release regions at different locations along the coast of the Gulf of Maine; they are the same size everywhere else in the model. C) The final distribution after 600 generations of a species released at the location of the red star in Long Island Sound, averaged over 1000 trials. D) Same as C, but for a release in Trinidad and Tobago. An interactive and zoomable version of (A) can be found at https://jamiepringle.github.io/PringleLushByers_InvasibilityInRealOcean/.

In the runs presented below, dispersal statistics are computed for larvae fixed to 1m depth and released during the months of April, May and June (the oceanographic northern hemisphere spring, when many Northern Hemisphere larvae are released) for the years 2007 to 2023. In the different runs, different species can, but do not always have, a difference in relative competitiveness expressed by a difference in reproductive output. At each location, 8 or more larvae are released per adult (depending on the model run and the relative competitiveness of the different species) and are allowed to drift for 22 days. Any that have returned to the habitat are eligible to settle. Those that have not, die. At a large majority of habitat patches, this number of larvae is sufficient to keep the population at or near the community carrying capacity.

In the online supplement, we show how to run these model results for different habitats, larval vertical locations, durations and timing, so different combinations of parameters can be examined.

### Results Part 1: Idealized model, Qualitative results and the importance of divergence

Using an idealized, linear and finite coast with various patterns of larval dispersal, the population model is used to find which patterns of alongshore variation in the magnitude and direction of larval transport leads to regions where competitively neutral newly introduced species are more likely to persist. The idealized model runs are initialized with a unique species at each location, all species are competitively neutral, and the model is run until stochastic population dynamics drive all but one or two species to extinction. These stochastic dynamics have been extensively studied in the literature of genetic drift, and the analogy between genetic drift in a haploid species and species frequency evolution in neutral populations has been well studied (e.g. Hubbell 2001; Vuilleumier and Possingham 2006 and citations therein).

When all species in the model are competitively neutral, the initial condition of a different species at each location is equivalent to a separate species being introduced into each location. The model is run several thousand times to understand how likely a species starting at any point in the domain is likely to persist – this is necessary because of the stochasticity of both dispersal and settlement success. The frequency with which a species introduced to a particular location eventually occupies the entire domain (hereafter “grows to fixation”) is a direct measure of how invasible a location is.

If there is no mean downstream dispersal of larvae (L_adv_=0) the stochastic dispersal will move the offspring of an adult uniformly throughout the domain, and the population dynamics will be roughly the same as a single well mixed population. The location of the species which grows to fixation is randomly spread throughout the domain (Figure **1**A), with a slight reduction near the edges of the domain (similar to Wilkins and Wakeley 2002). Since species introduced to all locations are equal, the resulting likelihood of fixation for a single introduced individual is roughly 1/N, where N is the total population of the domain (Kimura and Ohta 1969; Hubbell 2001). The median time to fixation in generations scales as the total population, though there is a long tail of fixations times (Figure **1**B and Hubbell (2001)). These results are consistent with an extensive literature of neutral population dynamics in closed populations.

Stochastic flows with no preferred directionality are, however, rare in the coastal ocean (Brink 2005). Directional flow will move larvae preferentially from one part of the domain downstream to another. This can make the offspring of otherwise identical species in one part of the domain more likely to survive than a species which starts in another part of the domain. When, in addition to the stochastic dispersal of larvae a distance L_diff_, there is a mean transport of larvae a distance L_adv_ downstream, the location of the ancestors of any given individual are more likely to be upstream of the individual’s location than downstream (assuming that *L*_adv_ is everywhere the same within the domain). Over time, the location of all the ancestors of individuals within the domain will be concentrated in a small region at the upstream edge of the domain. The size of this region has been shown to scale as L^2^ /L_adv_ (Wares and Pringle 2008; Teller and Pringle 2023). Because eventually all individuals in the domain will have ancestors which originated in this upstream edge region, individuals starting in or introduced into this region are far more likely to grow to fixation throughout the domain. This is seen in Figure **1**A for the case of a unidirectional mean flow, where the locations of the individuals which grow to fixation are seen to be concentrated at the upstream (left-most) portion of the domain. This region will be called the “Lineage Source Region” or “LSR,” because over time in this neutral case, all individuals in the domain will have descended from an individual in the LSR. The only way a species introduced outside of an LSR is likely to persist is to have a sufficient competitive advantage to spread upstream against the mean downstream flow (the magnitude of the competitive advantage needed to persist is the described in Byers and Pringle (2024)).

When the upstream LSR of size L^2^ /L_adv_ is smaller than the size of the entire domain, then the LSR’s population will be smaller than the entire domain. This will cause individuals in it to grow to fixation in the LSR much more quickly than they would if L_adv_ were zero (Figure **1**B and C), and the population of this upstream region will replace the total population size as the population which governs time to fixation, the stochastic extinction of other species, and other similar population dynamics (Wares and Pringle 2008; Teller and Pringle 2023). In analogy to similar terminology in population genetics, we call the population of this L^2^ /L_adv_sized region N_e_, the “effective population size.” As a single species in the upstream LSR grows to fixation, its offspring will spread downstream and become dominant in the rest of the domain (Wares and Pringle 2008).

The LSR formed by the unidirectional flow can be thought of as a region of divergent transport of larvae at the upstream edge of the domain. There will be no flux of larvae entering the upstream edge of the domain, since there are no adults that exist upstream of the domain, and thus no larvae released from outside of the domain. However, there will be larvae released in the LSR at the upstream side of the domain – and many of these larvae will be swept downstream of the LSR by the mean currents. The LSR is thus a location of a divergent larval flux, with no larvae entering the LSR while at the same time they are leaving the LSR.

To examine the case where the mean transport of larvae varies along the coast, two more models are run. In the first, the mean currents diverge, analogous to the divergence seen, for example, at the Mississippi River delta in Figure **2**A. In the second, the currents converge, as is seen near Cape Hatteras in Figure **2**A. The direction of larval transport is shown in Figure **1**A.

In the case with diverging currents, there is an LSR at the center of the domain (Figure **1**A) because it is from this region that larvae are exported, and so it is individuals which fixate here that go on to become dominant everywhere in the domain. Because the currents move in both directions away from the center, this region is twice the size as the region in the case where the current flows unidirectionally across the domain (Figure **1**A), and the time to fixation is about twice as large (Figure 2B). Nonetheless, the dynamics are similar to the case in which the mean flow enters through one boundary – the region which exports more larvae than it imports becomes an LSR, and individuals in these regions are very much more likely to grow to fixation. This is what one would expect from source/sink or metapopulation theory (Kawecki and Holt 2002; Kawecki 2004), where species in source regions are more likely to avoid extinction than those in sink regions.

Conversely, when the currents converge at the center of the domain (as currents converge at Cape Hatteras) there is no tendency for species originally at the converging region in the center of the domain to persist. The center is not an LSR. It is a region where a propagule rain falls from both sides, and so is a sink region for larvae originating on either side of the center. The species which persist are those which start near the two upstream edges where the flow enters, just as in the case of the upstream most edge in the case with unidirectional mean currents. Since larvae are moved from both edges of the domain to center, it is very difficult for a species which has grown to fixation on the left side of the domain to have offspring which reach the LSR on the right side of the domain and vice versa. Thus, instead of one species going to fixation everywhere, two species grow to dominate – one on the left side and one on the right side, with a species boundary at the location where the currents converge at the center of the domain (e.g. Pappalardo et al. 2015; Krumhansl et al. 2023). This can be seen in the time-history of the number of surviving species, in Figure **1**C, and is consistent with prior work on the origin of current mediated species range boundaries (e.g. Gaylord and Gaines 2000; Pringle et al. 2017).

In summary, neutral species which are introduced into regions which are net exporters of larvae are more likely to persist and grow to fixation than species that are downstream of the export, as one would expect from source/sink metapopulation dynamics. The LSR’s form where there is a divergence of the flux of larvae, either where the flow enters from outside the model domain, or where the currents diverge within the domain.

### Results Part 2: Oceanographically realistic models, Qualitative results and the distribution of LSRs

Using oceanographically realistic estimates of larval dispersal for a 22 day larval duration, coastal habitat and northern hemisphere spring larval release timing described above, locations where neutral introduced non-native species are more likely to persist along the coast of the Americas are identified and their origin qualitatively explained.

To capture the complexity of actual coastlines and oceanic dispersal, Lagrangian pathways from a high-resolution ocean model are used to quantify the connectivity between 1/12^th^ of a degree habitat patches in a habitat which consists of the 1/6^th^ of ocean nearest the coast. For computational efficiency, the habitat is broken up into regions roughly 25 patches in size (Figure **3**B). To quantify which regions are most invasible, 10 individuals of a novel neutral species are introduced into randomly chosen points in each region. The maximum population per patch is *P_max_*=1, and the model is run until the introduced species goes extinct or persists for greater than 600 generations. This is repeated 1000 times for each patch. The percentages of the introductions that have persisted is shown for the entire domain in Figure **3**A, and a subset of regions is shown in Figures **2** and **3**. The locations where fewer than 1% of the introductions persist are shown in light grey in Figures **3**. An interactive online figure is given in the digital supplement (https://jamiepringle.github.io/PringleLushByers_InvasibilityInRealOcean/) which shows both the fraction that persist along with the mean along transport of larvae (The arrows from Figure **2**) in a figure that can be enlarged to show any region. With a thousand random trials, an estimated persistence percentage of 1% has a 95% confidence interval of 0.5% to 1.8%, and an estimate of 0.1% has a confidence interval of 2.5×10^-3^% to 0.5% (Clopper and Pearson 1934).

In much of the coastal habitat, none of the introduced species persist, while only a very small fraction of introductions persist in others. Regions where greater than 1% of the introductions persist are relatively sparse and spatially isolated. Most species introduced in other regions go extinct in tens to hundreds of generations. As described in more detail in Byers & Pringle (2006), Wares and Pringle (2008) and Pringle et al. (2017), in these regions the larvae produced by adults settle on average downstream of the adults (where downstream is defined by the direction of mean larval transport). The adults are in turn replaced by larvae that originated upstream of their position, so that in time they become extinct at the point of introduction and eventually everywhere, unless at least one of their offspring settles in an LSR.

The locations where introduced species are more likely to persist follow the pattern described in the idealized models – they are locations from which more larvae are exported than imported, and so are LSRs. For example, the coastal region around the mouth of the Mississippi River Delta is one location where species are more likely to persist (Figure **2**). The mean transport of larvae is divergent there; the flow to the west of the region is towards the west, and the flow to the east is towards the east. As with the idealized results shown above, species introduced into this divergence region are much more likely to persist than those introduced elsewhere, because larvae are preferentially moved from this region to settle elsewhere, while few larvae from other regions are moved against the mean current by the stochastic transport into this region. Of course, the total reproductive output in the region, and the retention of some larvae within this region, must be sufficient to maintain the population in this region – the required magnitude of this reproductive output is the main focus of Byers and Pringle (2006) and in these model runs it is sufficient.

Introduced species are also more likely to persist when they start in large estuaries. Introductions into the Chesapeake and St. Lawrence are much more likely to persist than those introduced into other locations nearby (Figures **2**B and **2**C). The dynamics that lead to this persistence are the same as those described above. The mean transport of surface trapped larvae, the red arrows in the figures, is out of these estuaries. In the model, and for most species that can survive in an oceanic environment, there is no transport of larvae into the estuary from the river that feeds it. This leads to a divergence in the transport of larvae – none enter through the river, but some leave into the sea, making the estuary into a source of larvae to the coast outside the estuary and an LSR. As above, this assumes that the retention of some larvae within the estuary is sufficient to maintain the population within the estuary. Perhaps more importantly, as will be described below, this estuarine result is sensitive to the vertical behavior of the larvae.

A final region which would appear to be easy to invade in these models arises at the boundary between where the habitat is populated and where it is un-populated. Un-populated regions arise where there is enough loss of larvae out of the habitat that the larval production cannot sustain the population. An example for the larval duration, production and timing shown in Figure **3** is the Peruvian Coast. Figure **4** is a subset of Figure **3** for the west coast of South America, but with red dots illustrating where the potential habitat is occupied and arrows illustrating the direction of the alongshore current. There is a gap in habitation around Peru where the upwelling circulation moves most of the surface larvae offshore so that they fail to return to habitat. At the northern boundary of this empty habitat, along the coast of Ecuador, there is a LSR with a high likelihood of introduced species persisting.

**Figure 4:**
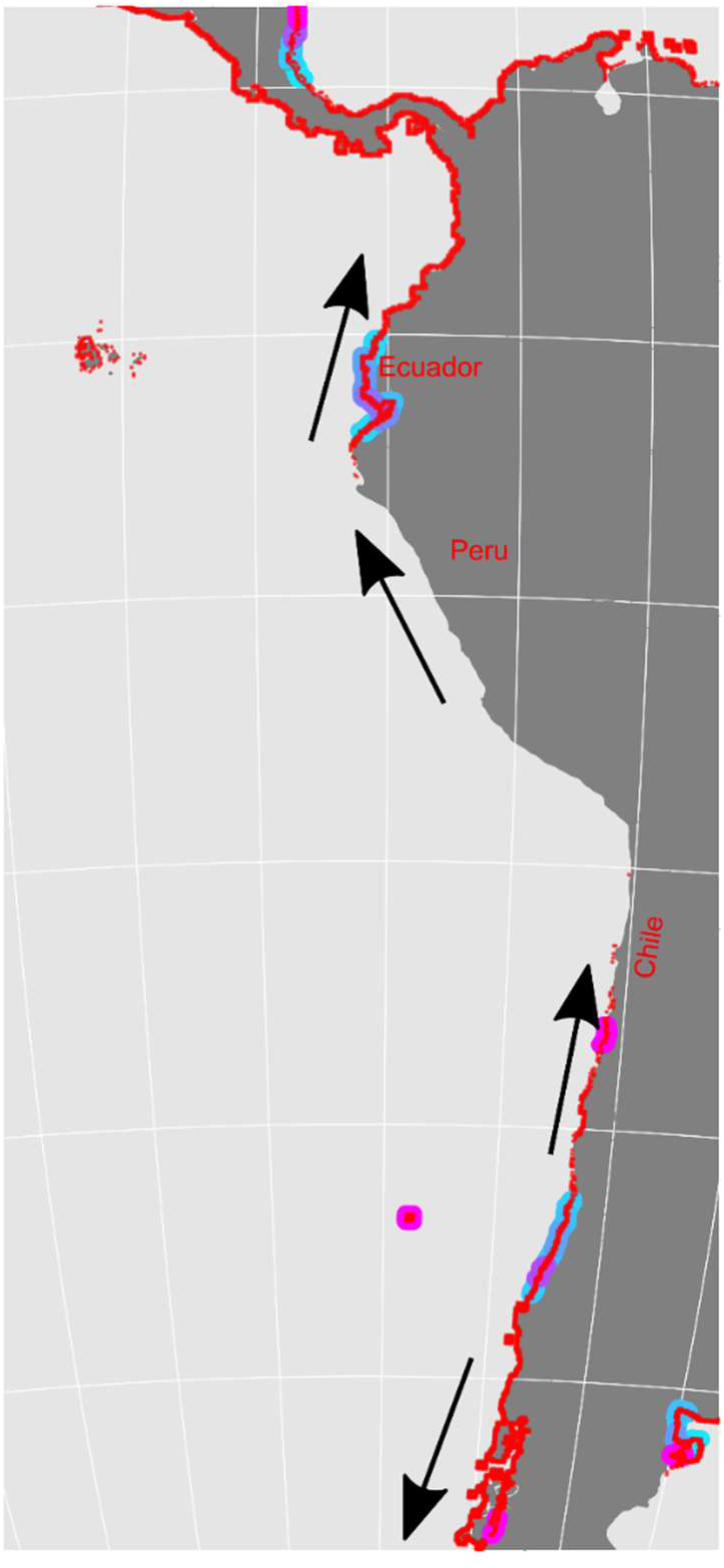
Same as Figure 3A, but showing as red dots all habitat patches which are occupied, and not showing habits in which < 1% of introductions are successful. The arrows show the alongshore direction of the geostrophic surface as described by Strub et al. (2019), Figure 1.

The origin of this LSR is straightforward – it is exactly like the idealized model result discussed for the unidirectional larval transport above. There is no flux of larvae coming from the uninhabited regions, and so the edge of the habituated region has a divergence of larval flux and becomes an LSR. However, this type of LSRs is likely an artifact of the model formulation. The model assumes the ecosystem consists of a large number of species with identical larval dispersal characteristics. The uninhabited regions are formed when the ocean circulation prevents any larvae from settling in those regions – typically, for these surface larvae, in regions of strong upwelling where the circulation moves surface larvae away from the coast. It is, however, highly unlikely that the existing species in a region would have a larval dispersal behavior that would minimize their reproductive success in that region. Modeling of species in upwelling regions show that larval behaviors that minimize larval loss will be strongly selected for (Pringle et al. 2014), and observations of larval behaviors in upwelling regions clearly show larval vertical behaviors which reduce loss offshore (Morgan et al. 2009; Morgan 2014). The uninhabited areas of the model are not uninhabited in reality, and the dominant dispersal behaviors of successful species in these regions do not match those in the model used to make Figures **3** and **4**. Instead, these boundary regions are likely to be locations where the optimal larval behavior shifts and thus are likely to be biogeographical transition regions between species with different larval behaviors, as described by Lush (2025).

In many locations along the coast, there is a low (but nonzero) likelihood of successful introduction over broad stretches of the coast – these are shown as the grey dots in **3**A, which indicate likelihood of persistence >0% but less than 1%. Most of these are in locations with no obvious flow feature which would allow an LSR to form. In most cases, introductions of a species into these regions lead to infrequent persistence when the descendants of the introduction drift downstream and are very occasionally able to enter an LSR and become established there. While LSRs are regions where there is much more larval emigration than immigration, there is sufficient variability in the dispersal that there are no locations that are entirely isolated from the coastal regions around them. For example, in figure **3**D, the distribution of a species introduced at the red star in Trinidad and Tobago is shown after 600 generations. The introduced species is not found at the point of introduction – instead, the descendants of the introduction moved downstream to an LSR in Jamaica and became established there and downstream (the currents can be seen in the interactive online figure). We cannot always assume that where an introduced species becomes abundant is where it was introduced.

Once an introduced species has established in an LSR, the offspring from that region will spread far downstream and become present in all the downstream sink regions. In Figure **3**C, the eventual distribution of a species introduced in Long Island Sound is shown. This distribution spreads from Cape Cod downstream to Cape Hatteras, and this distribution is established within tens of generations of the introduction. These geographical limits are set by ocean currents and the presence of other LSR regions, as has been documented observationally in this region (Pappalardo et al. 2015; Pringle et al. 2017). Similar dynamics have been shown to set biogeographic boundaries globally (Krumhansl et al. 2023; Lush et al. 2026).

### Results Part 3: Quantitative results from realistic modeling

The results presented above provide a qualitative picture of which regions are more susceptible to the introduction of a competitively neutral novel species, for one specific set of larval depth behavior, planktonic duration and release timing. However, they do not provide a quantitative framework for understanding how the invasibility of an area will change with the invasive propagule pressure, the local population density, or the relative competitive advantage of the invader with respect to local species. Further, the low population densities in the model results shown above will likely be far less than realistic population densities for many species – especially those which are habitat limited. It would be computationally infeasible to run the models with population densities appropriate to, for example, barnacles in the intertidal.

In this section, the correspondence of the results of our models which incorporate realistic dispersal to the well understood model of species dynamics in a closed well mixed population will be presented, and then these results will be used to quantify invasibility as a function of propagule pressure, competitive advantages of the introduced species, and habitat carrying capacity.

In an isolated population of size N where reproduction is asexual and offspring in a generation are chosen randomly from the adults in the last generation, the likelihood of a single individual’s offspring growing to a population N and displacing all other species is 1/N (Hubbell 2001). Thus if *n_intro_* individuals of a novel competitively neutral species are introduced into the isolated habitat, there is a

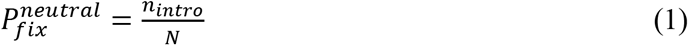

probability that they will eventually grow to fixation in the population. If initially every individual in the isolated population is of a different species, the mean number of generations until one grows to fixation will be approximately the same as the population, so N generations for a population of N individuals. These results, and a good summary of these population dynamics, can be found in Hubbell (2001), and they are consistent with the results for the species with no mean directionality of dispersal in Figure **1**.

We hypothesize that we can treat an LSR as if it were like the isolated population discussed above. If this hypothesis is true, we can estimate an effective population size N_e_ for the LSR for it by observing the likelihood of growing to fixation of *n*_intro_ larvae of single species introduced into the LSR *P*^neutral^ and rearranging equation (1) to give

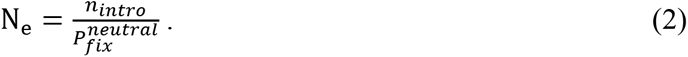

As in the genetic literature, N_e_ may not be the actual census population of the LSR – in particular, the boundaries of the LSR will not be sharp discontinuities, but will be fuzzy around the edges. For example, see the likelihood of fixation in Figure **1**A for the cases with mean larval transport. These likelihoods smoothly transition from some maximum to zero. N_e_ is a number that allows the dynamics of the LSR to be mapped to those of a closed population (Wares and Pringle 2008).

To test the hypothesis that the dynamics of the LSR can be understood as a closed population of size N_e_, a series of properties of the LSR will be tested and compared against the results which would be expected from this assumption.

From equation (1), it is seen that the fraction of introductions that grow to fixation should scale linearly with the number of individuals of the novel species introduced into the region. To test this, the model shown in Figure **3**A is repeated with an *n*_intro_ of 1, 5 and 10 individuals, and the probability of fixation is shown for each as a function of the probability of fixation after the introduction of a single individual. As is expected, there is a five-time greater likelihood of fixation when 5 individuals are introduced and a 10 times greater likelihood when 10 individuals are introduced (Figure **5**A). This is consistent with the model. (The larval production R is 8 for all models and *P_max_*=1. The correlations of the log-transformed data between the expected and observed fixation rates for *n_intro_* of 5 and 10 are 0.94 and 0.92, and these are significant at P>0.05.).

**Figure 5:**
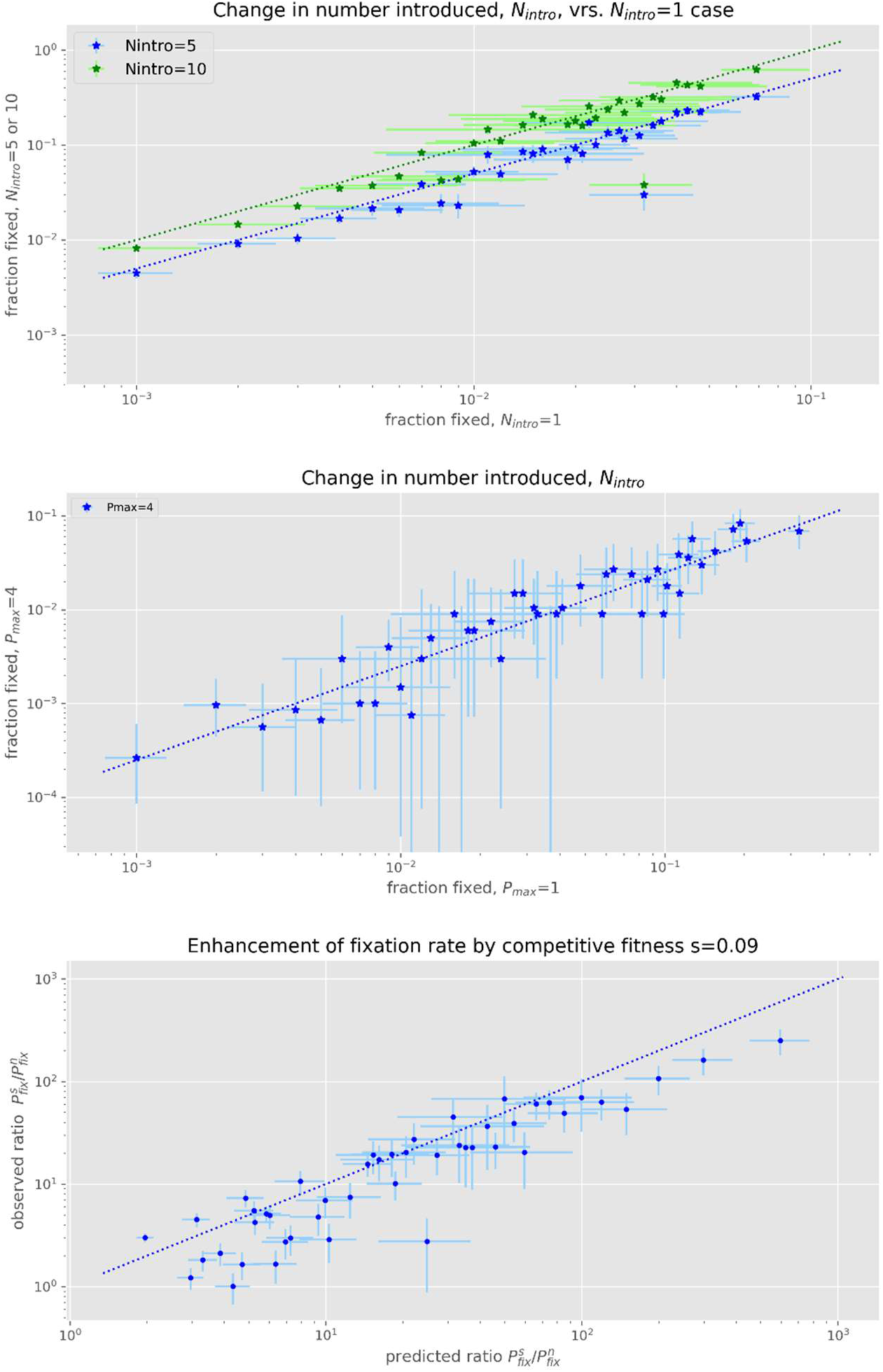
A) fraction of introductions which persist for 600 generations with 5 or 10 introduced individuals as a function of the fraction which persist for 1 introduced individual. Here and below, the lines are the expected values derived from a closed well-mixed population, and the error bars are 95% confidence intervals. B) Fraction of introductions which go to persistence after the introduction of 5 individuals into a habitat with a carrying capacity of *P_max_* 1 (horizontal axis) or 4 (vertical axis). The line is the expected 1:4 line. C) ratio of the likelihood of fixation for a species with a 10% reproductive advantage (*s=0.09*) to a neutral introduction for 1 introduced individual and a carrying capacity of *P_max_*=1. The vertical axis is the observed ratio, the horizontal is the ratio predicted by equation 6, and the line is the 1:1 line.

The likelihood that that an introduction will grow to fixation will depend on the population density – if the analogy to a closed populations holds true, when there are twice as many adults per population patch, then the total population of a region will be twice as large, and the likelihood of fixation for a new species with a given rate of introduction will be half as big (equation 1). In Figure 4B the likelihood of the introduced species growing to fixation for a habitat with carrying capacity of 4 adults per species is compared to one with a carrying capacity of 1 adult per species, and as expected, the likelihood of fixation is a quarter as large. (R=8. The correlation of the log-transformed data is 0.94 and is significant at P>0.05.) Since real population densities in the ocean are almost certainly different than those in the model, the actual N_e_ in the ocean will be related to that observed in the model by the ratio of their population densities H_dens_ per area:

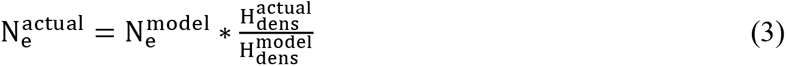

Where 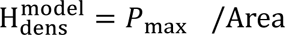, where Area is the size of a patch 1/12^th^ of a degree on a side, 86km^2^ * cos (latitude) squared kilometers.

The models presented so far have assumed all introductions are competitively neutral. This is highly unlikely to be true for actual introductions of non-native species (Byers 2009). The effects of differences in competitiveness between the introduced and native species are analogous to the effects of selection on a haploid allele in a closed population. Following Kimura and Ohta (1969) but for a haploid population, the likelihood of growth to fixation of an introduced species with a relative competitive advantage *s* is

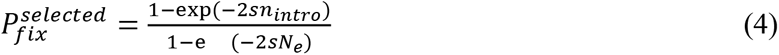

Where s is defined as the competitive advantage of the introduced species with respect to the native species:

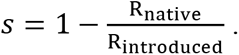

R_native_ and R_introduced_ are measures of the potential reproductive success of the native and introduced species, respectively. They are the number of larvae that are released by each adult which would survive dispersal, settlement, and post-settlement mortality until they are competent to reproduce if all larvae reached suitable habitat. It is assumed that the native and introduced species have the same dispersal. The fractional increase in likelihood of growing to fixation with selection is then

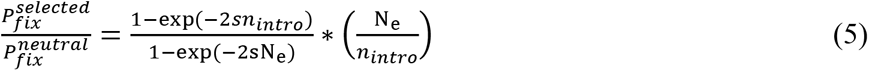

These equations can be tested by making the growth R_invasive_ for the introduced species 10% larger than that for the native species (8.8 and 8.0 respectively), so that *s*=0.09, and comparing the relative rate of fixation in each LSR to that in the neutral model. Figure **5**C compares 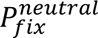 against the relative increase of the probability of growth to fixation between these two models, 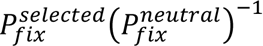, and the comparison is good, with a significant correlation of 0.92 for the log-transformed ratios.

All the results shown in Figure **5** support the hypothesis that the population dynamics of the LSR regions can be understood by thinking of them as a closed population of size N_e_ as calculated with equations (2) and (3). This is not to say they are a closed population – they export larvae to downstream regions. But N_e_ can be used to estimate the invasibility of these regions as a function of propagule pressure, habitat density, and relative species competitiveness.

### Discussion: Alongshore variation in larval transport can make it easier for non-native species to persist

In Byers and Pringle (2024), it was argued that for open coastlines with little variation in the alongshore transport of larvae, the mean downstream transport of larvae would make it much more difficult for introduced non-native species to persist. The population model they used had a different structure than the one here – in particular, their model was continuous, instead of having discrete individuals. This means that instead of giving a likelihood that a countable number of introduced species will persist, it gives a criterion for when any introduced species will persist – essentially, it assumes the carrying capacity is everywhere very large. In the appendix, Byers and Pringle (2024) is translated into the notation of this paper. They found that the introduced species would need a competitive advantage *s* of

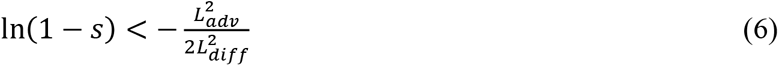

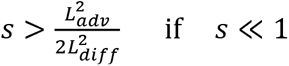

to persist. *L_adv_* is the mean downstream distance a larva is transported, and *L_diff_* is the standard deviation of the transport distance of all the larvae. Equation (6) shows that if the mean downstream transport distance *L_adv_* is greater than zero, than it will take a finite *s*>0 for the introduced species to persist, in the limit that the dispersal parameters are approximately alongshore uniform. A quantitative discussion of the meaning of “alongshore uniform” is given in Byers and Pringle (2024). To a good approximation, if equation (6) is satisfied by an introduced species over an alongshore distance large compared to *L_adv_* and *L_diff_*, then the introduced species is very likely to persist.

In Figure **6**A, the critical *s* that must be exceeded for an introduced species to persist under the assumptions of Byers and Pringle (2024) is shown for the larval parameters used in this manuscript (April-June release, larvae fixed to the surface, 22 day planktonic duration). Colors are only shown where the critical *s* is equal or greater to 0.1. Almost all of the coast is colored, indicating that for nearly all the coast Byers and Pringle (2024) suggests a critical *s*>0.1 is needed for an introduced species to persist.

**Figure 6:**
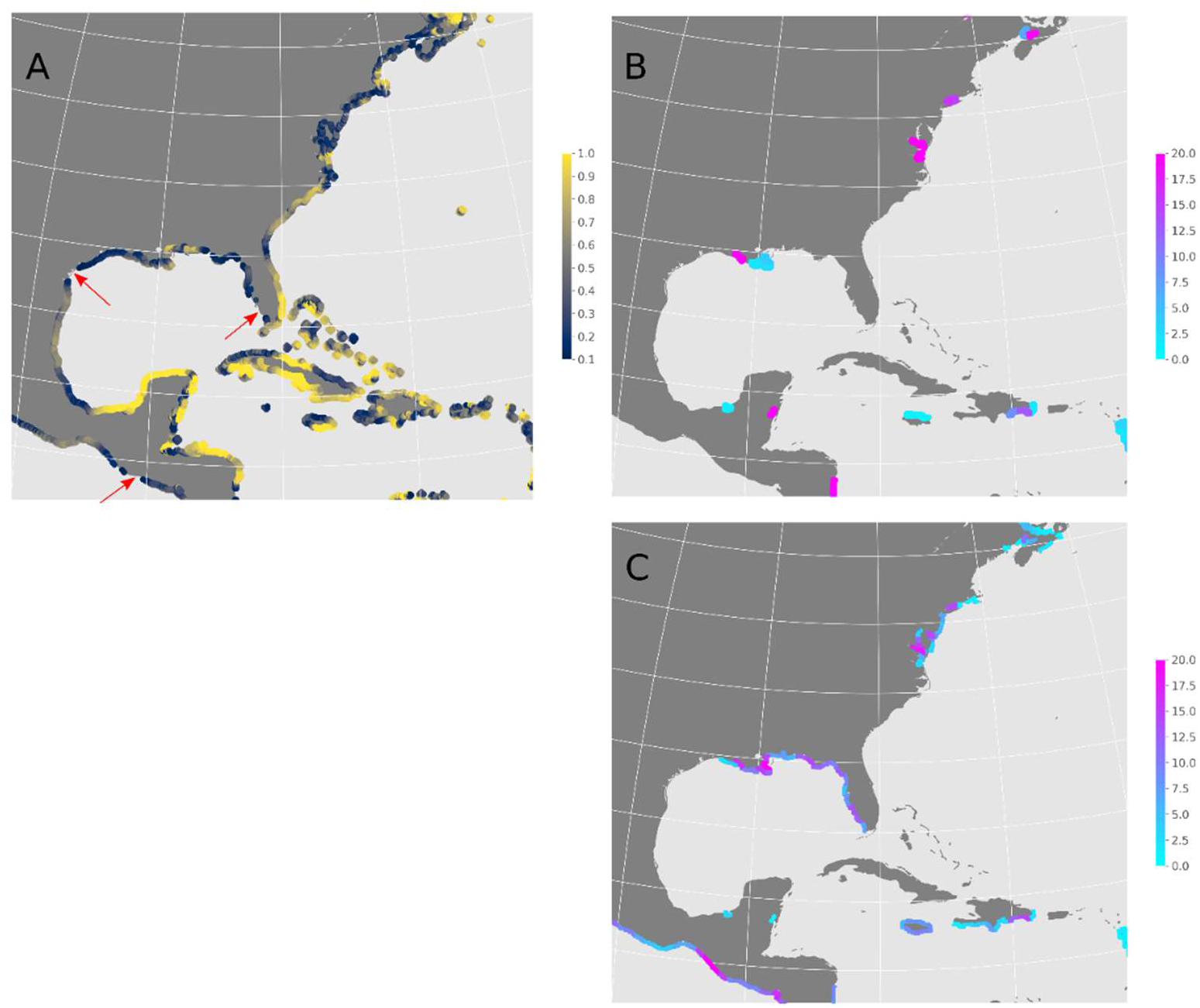
A) the critical competitive advantage *s* needed for a species to persist under the assumption that there is no spatial structure in the currents, from Byers and Pringle (2024) and the appendix. The critical *s* is only shown where it exceeds 0.1. Red arrows indicate regions where the critical *s* is less than 0.1. B) The fraction of neutral introductions which persist for the introduction of 10 individuals for a model with a carrying capacity *P_max_*=1, as calculated for 1000 introductions. C) The same as B) but the introduced species has a competitiveness advantage of *s*=0.09 (equivalent to a 10% greater larval production than native species).

Figure **6**B shows the fraction of introductions of a neutral (*s*=0) species that persist when introduced at different locations in this region. Along most of the coastline, there is no persistence – it is only in some isolated regions that there is any likelihood of persistence. These are mostly associated with regions of divergent larval transport (Figure **2** and online interactive figures) or estuaries. These are the LSRs discussed above. Because they are sources of larvae, once the introduced species becomes established in the LSR, the larvae that are exported from the LSR will spread its range until limited by other dynamics (Figure **3**C and Lush et al. (2026)).

If the introduced species has a greater reproductive potential than the native species, we expect it to be much more likely to persist (equations (5) and following). Introductions would be expected to persist when they are either into a location which is an LSR for neutral introductions or if they are introduced into a region where their competitive advantage is sufficient to satisfy equation (6) from Byers and Pringle (2024). In Figure **6**C, the fraction of introductions which persist are shown for an introduced species with a 10% greater reproductive output than the native species (*s*=0.09) – this is just less than the threshold for displaying critical *s* in Figure **6**A.

As would be expected, all the LSRs seen in Figure **6**B are locations where the selectively favored species has a high likelihood of persistence. Most locations in Figure **6**A that are uncolored because equation (6) suggests that they can be settled by a species with an 0 ≤ *s* ≤ 0.1, such as part of the west coast of Florida and the coast of Guatemala (marked with red arrows in the figure), are also regions where introduced species has a high rate of persistence. This is what would be expected from Byers and Pringle (2024).

In addition, there is some persistence of introductions of the competitively advantaged species into the regions adjacent to the LSRs shown in figure **6**B and adjacent to the regions where Byers and Pringle (2024) suggest that these introductions would persist. There could be two reasons for this. First, the descendants of the introduced species could spread to adjacent LSRs and establish there. Or there could be regions such as the Delaware estuary which have too much in-migration of larvae to be an LSR – but still have less in-migration of larvae than a location on the open coast. If the in-migration rate is less than the competitive advantage *s* of the introduced species, then the introduced species could become favored to persist. This is the classical metapopulation result that a species can persist in the face of immigration of larvae of competing species if its competitive advantage is greater than the immigration rate (Hubbell 2001). Which of these two mechanisms is more important is an interesting question for future research.

Alongshore circulation patterns can also, however, make some regions more difficult for introduced species to persist in, even if the species have a competitive advantage. Near Corpus Christi TX, in the northwest corner of the Gulf of Mexico, Figure **6**A shows a location where Byers and Pringle (2024) would predict persistence for an introduced species (marked by a red arrow). But in neither Figure **6**B or **6**C is there any indication of neutral or selectively favored species persisting there. This is because, as shown in Figure **2**, the larval transport is convergent at this location, and as discussed above and shown in Figure **1**A, regions of convergent flow are highly unlikely to allow an introduced species to persist.

### Discussion: The importance of a competitive advantage as population density increases

In the simulation presented above, the stochastic dynamics of a finite population have played an important role. This has been helpful, for it allowed straightforward tests of the hypothesis that the LSR could be treated as a population of size N_e_. But the population maximum per patch *P_max_* in these models of 1 to 10 individuals per 1/12^th^ of a degree patch (≈ 60km^2^ at mid-latitudes) is very much less than one would expect for many species.

It is clear from Equations (1) and (3) that as the population density of the LSR increases, the native population of the LSR increases and the likelihood of persistence of a fixed number of introductions of a neutral non-native species becomes very small. Equations (5) and (1) can be combined to show that in the limit that 2*sn*_intro_ « 1, the likelihood of persistence and successful introduction also becomes very small, of order *n*_intro_/*N*_e_. However, as the relative competitive advantage of the introduced non-native species increases, this is no longer true. If the number of introduced individuals and their advantage *s* grows large enough that 2*sn*_intro_ > 1, and we assume that the number introduced is smaller than the effective population size, than both exp (−2*sn*_intro_) and exp (−2*sN*_e_) become small, and from Equation (4) we see that the likelihood of the introduced species reaching fixation goes to 1. So if 2*sn*_intro_ > 1 in an LSR, the likelihood of the introduced non-native species persisting in that LSR becomes close to 1. If, as seems likely, the number of non-native individuals introduced into a LSR is much less than the population of species it competes with for habitat in the region, then the non-native species must have a competitive advantage over the native species to be likely to persist.

This is consistent with what has often been assumed (Byers 2009), that stochastic population dynamics are unlikely to allow a neutral introduced non-native species to persist and that invaders must have a competitive advantage over native species to be likely to establish. It is straightforward to show, however, that the required competitive advantage will be much less in an LSR than on the open coast. In Pringle (2024), it is shown that L_adv_ and L_diff_ are often of roughly the same magnitude, so that Equation (6) predicts that *s* must be greater than ≈ 0.4 for a species introduced on the open coast to persist. However, the results above show that in an LSR *s* must only exceed ≈ 1/(2*n*_intro_) to persist, which will be less than 0.4 even if only two individuals are introduced, and will be much less if there are greater numbers of introduced non-native species.

## Discussion: Estuaries

Many estuaries are hot-spots for the appearance of non-native species, and more invasive species are found in estuaries than on the open coast (Preisler et al. 2009 and citations therein). This is often attributed to the greater number of anthropogenic introductions into estuaries which contain ports (Preisler et al. 2009). The results above suggest an additional and complementary mechanism. In Figures **2**, **3** and **6**, large estuaries such as the St. Lawrence River and the Chesapeake are seen to be LSRs, making them particularly vulnerable to the introduction of non-native species. However, this result is not general to all larval behaviors and is sensitive to the vertical distribution of the larvae with respect to the estuarine circulation.

Most estuaries are regions in which a relatively small amount of river water is added to a much larger volume of ocean water to produce an outflow which is somewhat fresher than the ambient ocean water, and much closer in salinity to the ocean than to that of the river (Geyer and MacCready 2014). In most estuaries with moderate tides (Hansen and Rattray 1965), the outflow is in the upper part of the water column as it is less dense, and the inflow is in the lower part of the water column. In the simulations above, the larvae were in the upper portion of the water column, and so were transported out of the estuary towards the sea. In these estuaries, this circulation leads to a divergence in the transport of larvae, since none enter the estuary from the river, and many leave the estuary to the ocean. However, if the larvae move to a deeper depth, the net flux of larvae will be from the coastal ocean into the estuary. This is equivalent to a convergent flow, as more larvae will enter than leave the estuary. The estuary is a sink, not a source, for larvae which remain deeper and will not to be an LSR for species with this larval vertical behavior.

## Discussion: Changing climate & Shifting ranges

It is often assumed, and has been well observed, that as the ocean’s climate changes, species ranges will shift as a result – usually poleward (Sunday et al. 2012, 2015; Pinsky et al. 2013). If the poleward shift of a species with planktonic dispersal is against a mean equatorward current, as seen in the US Mid-Atlantic Bight and poleward (Figure **2**), it will not occur simply because larvae are moved downstream by the mean currents. Instead, less frequent stochastic upstream transport events must introduce “non-native” pioneers into their potential new habitat.

These introductions will be more successful if they are introduced directly into an LSR – thus we hypothesize that the poleward shift of species will be driven by initial introductions into LSRs, and we will see sudden jumps in species ranges from their current range to the next poleward LSR. Once in the LSR, they will spread via larval transport into regions downstream of the LSR. This also suggests that the first place that poleward moving species will be seen in a region will also be places subject to increases in the number of anthropogenically introduced non-native species, since both the anthropogenically introduced individuals and the poleward moving migrants are more likely to persist if introduced into an LSR.

## Discussion: Limitations in these results originating from the circulation model

The estimates of larval dispersal which define the connectivity of the habitat and drive the population model, and the spatial habitat of the population itself, are defined on the underlying circulation model’s 1/12^th^ of a degree grid. The resolution of the underlying Mercator GLORYS global model ranges from 9 km at the equator to 5 km at 45° latitude to about 4.6 km at 60° latitude. This resolution, while at the cutting edge of global models, is too coarse to resolve many estuaries. Even the large St. Lawrence, Bay of Fundy and Chesapeake estuaries are only marginally resolved (see the grid points shown in Figures **2**B and C and **3**B and C). In addition, this model does not include tidal flows. So while the average circulation for these regions in the model are consistent with shelf flows in general (Wilkin and Hunter 2013; Amaya et al. 2023), and the mean circulations in the model match those observed in these estuaries and for the shelf LSRs we discuss (El Sabh (1976) Fig. 11 for The St. Lawrence, Mavor (1921) and Aretxabaleta & McGillicuddy (2008) for the Bay of Fundy, O’Donnell et al. (2014) and Vieira (2000) for Long Island Sound, DiMarco (2005) et al. Fig. 4 for the Mississippi outflow region), the fact remains that the model is not optimal for large estuaries and will not resolve estuaries smaller than the Chesapeake or St. Lawrence estuaries.

There are higher resolution but limited spatial domain models of estuaries and coasts (e.g. Chao et al. 2017, a 200m resolution model of San Francisco Bay; Bever et al. 2021, a 1km resolution model of the Chesapeake). But to be useful in the population modeling described here, larval dispersal must be defined over a domain that encompasses many LSRs, to avoid the corrupting effects of model boundaries – model boundaries across which there is an inflow will form an artificial LSR, for the same reasons as described for the unidirectional flow shown in Figure **1**. It should be possible to combine estimates of larval dispersal from highly resolved regional models with the results from larger scale models of the open coast – but has not yet been done and remains an area of future research.

Thus, while the work above identifies estuaries as potential LSRs for species with the appropriate vertical behavior of their larvae, the results shown in Figure **3** are likely to miss LSRs which form in smaller estuaries which are poorly or not resolved by the 1/12^th^ of a degree Mercator GLORYS model. Our results highlight the need to assemble a global estimate of connectivity which resolves smaller estuaries on a continental scale. This will become possible if Lagrangian particle models are used to combine circulation models from high-resolution regional models and larger scale, lower resolution models.

## Discussion: Limitations in this framework arising from the population model

The population model used above has focused on the role of circulation in creating regions that are susceptible to invasions (LSRs) because of the alongshore variation in larval transport, deliberately neglecting many other mechanisms that are present in the actual ocean. All results presented above assume the potential habitat of all the species in the model, both native and non-native, is the same everywhere and exists everywhere along the coast. The model also assumes that the introduced species, and all the species it competes with, share the same dispersal dynamics. While these assumptions are useful in isolating the role of ocean circulation on the invasibility of different locations along the coast, they neglect much that is important to actual inter-species interactions.

Perhaps most significant is the assumption that the introduced species and its native competition share the same dispersal dynamics. To understand how this assumption affects the potential ability of an introduced species to persist in an LSR, we can think of an LSR as a single well-mixed population in which some larval production is retained, and some is exported – as described above, this is a reasonable simplification. If we have two species in this population with equal reproductive output, larval mortality, competitive ability, and larval dispersal behavior, they will have equal competitiveness, and their relative population will only fluctuate due to stochastic population dynamics. But now imagine one species has slightly different dispersal behavior that leads to more retention of its larvae within the population and less migration out of the population. Presumably, that species’ population would grow and dominate the LSR. At a minimum, this suggests that the dynamics of dispersal within an LSR could alter the relative competitiveness of species within the LSR. The relative competitiveness implications of different dispersal dynamics have been examined in riverine and spatially uniform coastal environments (Lutscher et al. 2007; Pringle et al. 2014; Peniston and Burgess 2024) – but how differences in dispersal behavior interact with oceanographically realistic dispersal patterns to make certain areas easier to invade remains a rich space for future research.

The interaction of the vertical behavior of larvae with the cross-shelf circulation can lead to different species with different larval behaviors having very different rates of larvae returning to suitable habitat. This larval return rate will vary along the shelf as the cross-shelf current structure varies along the coast. Lush (2025) discusses how this can lead to species range boundaries along the coast, and Byers and Pringle (2024) discuss how these differences in vertical larval behavior can affect the ability of introduced species to persist.

It has also been assumed that all the species being considered have potential habitats that spread along the entire coastline. But of course, this will not be true for most species. If all species have a shared tolerance for habitat – for example, they all require a hard substrate which does not exist upstream of some location – and so have a shared spatial boundary to their habitat, the result is straightforward. If the flow enters from outside the habitat, it cannot bring larvae from outside of the habitat, but larvae released just downstream of the habitat boundary will move on average downstream, and there is a divergence in the flux of larvae across the boundary. This is the same as the dynamics at the edges of the idealized domain in Figure **1** for the unidirectional or converging mean flows. As in those cases, the habitat boundaries the flow crosses become LSRs. This can be seen in Figure **3** on the Labrador coastline at 60N. This is the northern extent of the model domain, and the flow across this boundary is into the domain. It thus becomes an LSR not because of the flow in this region, but because it is the edge of shared species habitat through which currents enter.

If, however, one species’ habitat ends at a location on the coast, and a neutral competing species’ habitat ends upstream of that location, then one would not expect an LSR to form at the habitat range boundary of the first species in the absence of any alongshore variation in the transport. The second species’ larvae would flow across the habitat boundary of the second species, and these larvae would be expected to reduce the population of the first species downstream of the boundary over time, until it goes extinct at its range boundary. These dynamics are described further in Pringle et al. (2017).

There are likely many other interactions between species’ habitats, dispersal behavior, phenology and the alongshore variation in coastal flows to be examined. The developments above should form a framework to begin this exploration.

### Text Box: Comparisons with observations of range expansion: *Carcinus maenas* in the Canadian Maratimes after the mid 1980s

We predict above that certain regions are more vulnerable to the introduction of non-native species because of the alongshore variation in larval dispersal. Confirming that this is true is not straightforward, for many of the locations we predict to be more invasible are also likely to have more introductions – e.g. many large estuaries we predict to be LSRs are also major ports, so there are two linked mechanisms which might explain the initial appearance of a novel species. However, the distributions of LSRs may be apparent when a species expands its range. We hypothesized above that when a species expands its range, it will be first observed in the locations most vulnerable to introduction – the LSRs. Subsequently the introduced species will spread out from the LSR, given suitable habitat. The timing of species range expansion can thus be compared to the prediction of the model above.

Below, the history of the spread of *Carcinus maenas* (Figure **7**) in the Canadian Maritimes northward of Halifax in the late 1980s to the present is compared to predictions made with the model above, and some support is found for the dynamics described above. In the accompanying online supplement, configuring the model for an arbitrary region, larval duration, larval vertical behavior and phenology are described using this system as an example.

**Figure 7.**
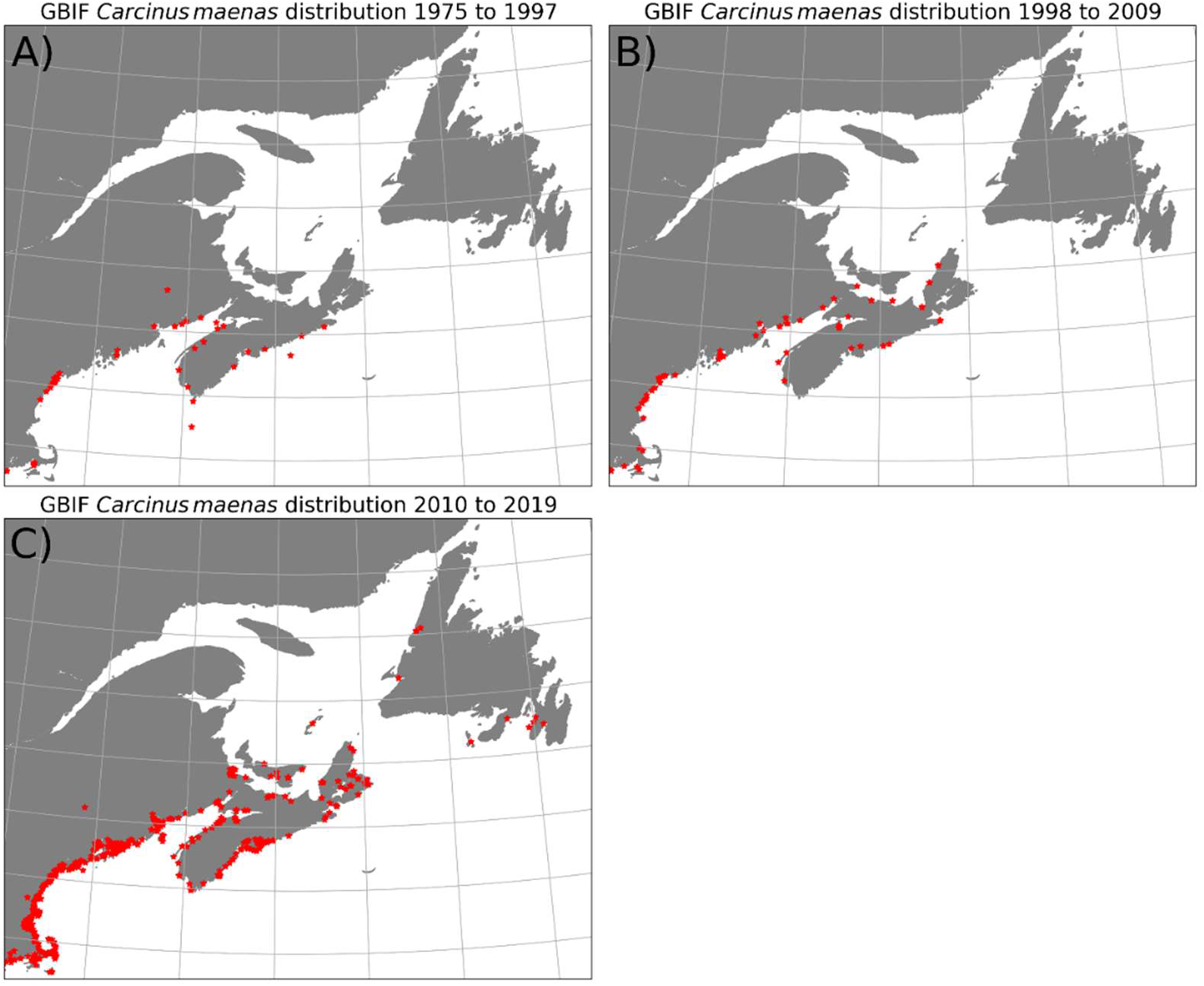
Global Biodiversity Information Facility (GBIF) occurrence data for *Carcinus maenas* for observations from A)1975 through 1997, B) 1998 through 2009 and C) 2010 through 2019.

*C. maenas* was introduced into Mid-Atlantic Bight from Europe in the 19^th^ century, and by 1960 had spread to Halifax NS (Audet et al. 2003) (All place names are shown in Figure **8**). Its range was thought to be limited by temperature, and on particular cold winters would disappear from the Western Gulf of Maine and Scotian Shelf (Welch 1968; Audet et al. 2003). In the late 1980s and early 1990s, there was an introduction from Europe of *C. maenas* into Cape Bretton Island and the Bras d’Or salt-water lakes; these individuals caried genes which allowed them to tolerate lower temperatures (Roman 2006; Jeffery et al. 2018). These cold-tolerant genes then allowed *C. maenas* to spread west to the Northumberland Strait and, by about 2002, north-west to Newfoundland (Audet et al. 2003; Blakeslee et al. 2010, Figure 8).

**Figure 8.**
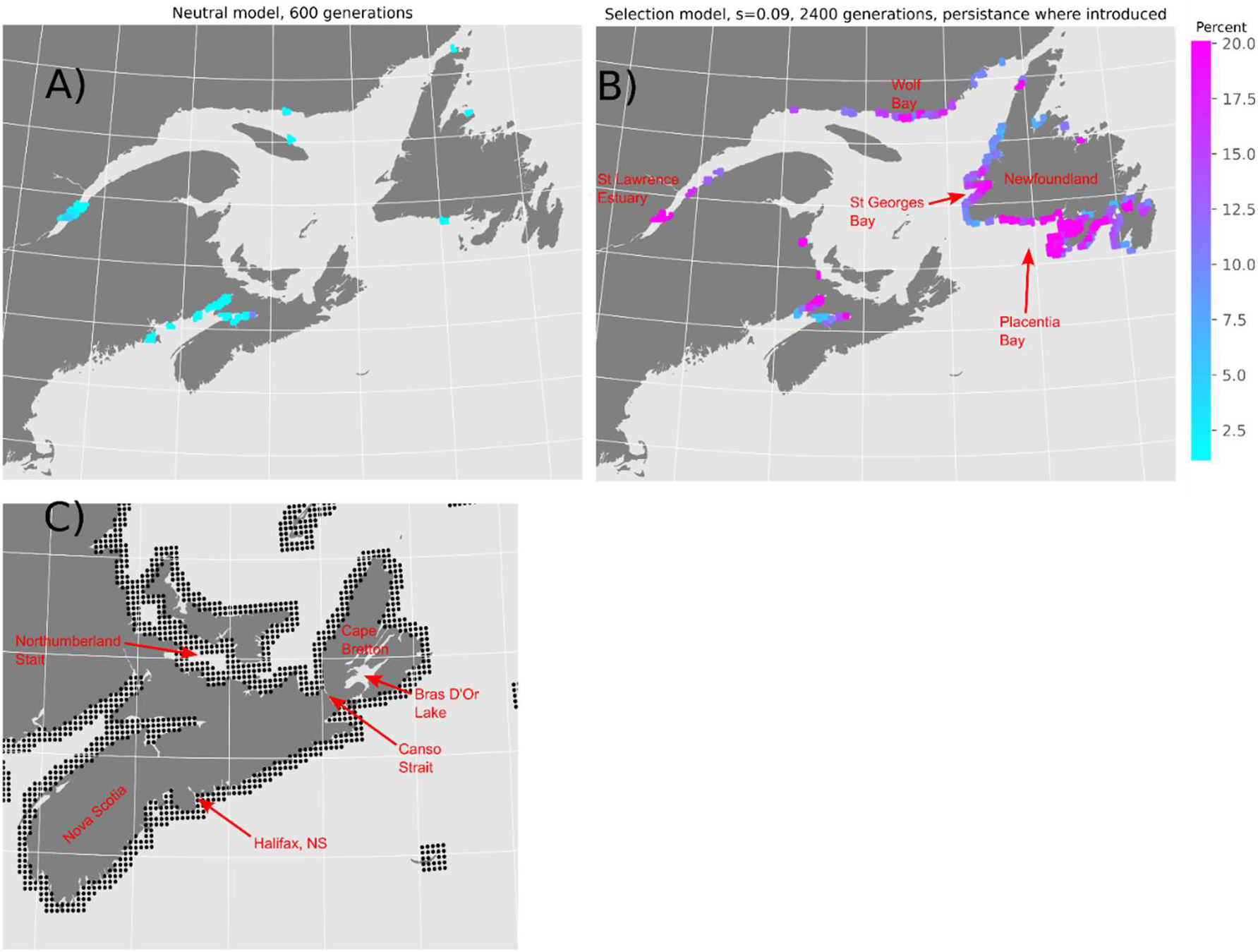
A) likelihood of persistence for a neutral introduction of 1 individual into a habitat with carrying capacity of 1 per patch. Persistence defined as the introduced species being present after 600 generations. B) Same as A, except the introduced species is favored by a 10% greater fecundity than the native species (*s*=0.09). Persistence only includes species that are still present where they were introduced. C) Black dots indicating habitat patches in model around Nova Scotia. Note that the model does not have connectivity through Canso Strait and does not resolve the Bras d’Or lakes.

To model *C. maenas*, we assume a habitat confined to within 1/6^th^ of a degree of the coast, a larval duration of 40 days and release in the climatological May and June calculated for the years 2007 to 2023 (Pringle et al. 2011 and references therein). The larvae are fixed to the surface, which is a simplification of larval depth behavior which has been seen in some locations in Europe to shift from the surface to greater depths as the larvae mature (Queiroga 1996); this model does not have the ability to simulate these ontogenetic shifts in larval behavior. In Figure 8A, the LSRs are shown for a neutral invader, and in panel B for an introduction of a selectively favored introduced species with an s=0.09; in this figure, only introductions which lead to finite populations at the region of introduction are shown.

The first post-1980 range expansion occurs with the introduction of a novel genotype of *C. maenas* into the Bras d’Or saltwater lakes, Canso Strait, or someplace nearby. The model does not identify these regions as a potential LSR, and so part of the hypothesis is failed. It is likely, though cannot be proven without further modeling with a higher spatial resolution, that this failure occurs because the 1/12^th^ degree Mercator model is too coarse to include either Canso Strait or the Bras d’Or lakes (Figure **8**C) – this failure mode is described above (see: Discussion: Limitations in these results originating from the circulation model).

Sometime around 2002, *C. maenas* appears in Placentia Bay, Newfoundland. Blakeslee et al. (2010) found it to first occur near oil terminals in upper reaches of the Bay (their figure 1), and found that the genetics of the introduction were consistent with anthropogenic transport of individuals from southern Nova Scotia, and not transport (natural or otherwise) from the more nearby Cape Breton Island populations. The modeling with a competitively favored introduced species (Figure **8**B) shows this region to be a potential LSR, consistent with our expectations, and subsequent observations have shown *C. maenas* to have expanded to the outer portion of Placentia bay and other nearby bays (Figure **8**C and (Government of Canada 2018)). By 2008 it was next identified in St. Georges Bay (Government of Canada 2018), which the model identified as the other most likely location for an LSR (Figure **8**), and has subsequently spread to more locations around Newfoundland (Figure **7**C).

The next adjacent regions that the model with selection and dispersal parameters appropriate to *C. maenas* predicts as LSRs are the north coast of the Gulf of St. Lawrence around Wolf Bay, Quebec and the St. Lawrence River, suggesting that *C. maenas* could persist and spread from them if introduced there. Given the closeness of the former to existing populations, and the extent of shipping to the latter, it will be interesting to see if *C. maenas* is soon observed in these locations.

On a larger scale, this comparison of observed and expected changes in the distribution of a cosmopolitan invader illustrates what is needed for better understanding and prediction of the spread of invasive organisms. It is important that we develop more highly resolved coastal models and couple them to larger regional models so we can estimate connectivity between small embayments and other coastal features over large length scales. It is important to have quantitatively accurate estimates of the vertical behavior and phenology of larvae. These can be used to estimate where invaders are more likely to persist when introduced. To test these predictions, we need routine surveying of the biogeography of the coastal ocean to accurately determine how the spatial distributions of species evolve.

## Discussion: Future directions

The spatial structure of LSRs and the locations to which descendants of the LSRs spread have important and testable implications for the spatial structure of genetic diversity within species with planktonic dispersal (Pringle and Wares 2007; Wares and Pringle 2008; Teller and Pringle 2023). In particular, LSRs are expected to generate strong, spatially localized signals of genetic structure despite open population connectivity. The interactions of LSRs with coastal circulation and life history traits to spatially structure genetic diversity in a species with planktonic dispersal represent a promising direction for future work and are the subject of ongoing investigation.

The model presented above is intentionally very simple and designed to isolate the effects of spatially structured dispersal. It assumes that all competing species have the same dispersal and phenology, their competition occurs solely through their neutral competition for (i.e., neutral occupation of) habitat, and their relative competitiveness is only expressed through different reproductive output. Relaxing any of these assumptions— including species-specific dispersal, stage-structured demography, non-neutral interactions, or environmentally dependent fitness— is expected to reveal additional mechanisms by which oceanography modulates competitive outcomes of coastal species with planktonic dispersal. Thus, this work represents a first step toward a more general theory of how spatially structured dispersal interacts with life-history to shape biogeography under climate change, range expansion, and anthropogenic species introductions.

## Significance Statement

The successful establishment of non-native marine species often depends on how planktonic larvae are transported by coastal ocean currents, yet the role of alongshore variation in these currents remains poorly understood. A long-standing assumption is that invasion success is greatest in highly retentive coastal regions, but there is no general framework for identifying which circulation features actually promote persistence after introduction. By combining large-scale ocean circulation models with open-source software we have developed, we show that regions with strong alongshore divergence in larval transport— rather than retention—are disproportionately susceptible to invasion because they function as effective larval source regions, providing a mechanistic link between coastal circulation, population persistence, and invasion risk that can be applied across species, coastlines, and changing environmental conditions.

## Breadth of interest

We provide a theoretical basis and software to quantify what parts of the global coastal ocean are most susceptible to the introduction of non-native species as a function of their dispersal behavior. We quantify the likelihood of a successful introduction as a function of local circulation and the relative fitness of the introduced species. It will be broadly useful to managers and ecologists interested in the spread of invasive species and the shift in species ranges with a changing climate.

If possible, we wish the section “Comparisons with observations of range expansion: *Carcinus maenas* in the Canadian Maratimes after the mid 1980s” to be printed as a text box

## Author Contribution Statement

**James M. Pringle:** Conceptualization, writing – original draft, software. **William G. Lush**: Conceptualization, writing – reviewing and editing. **James E. Byers**: Conceptualization, writing – reviewing and editing.

## Acknowledgments

This work was supported with NSF funding from OCE-1947954 and OCE-1947884. We are grateful for the feedback from John Wares on the manuscript and early presentations of the ideas within it.

## Appendix: Converting from the notation of Byers and Pringle (2024) to that of this work

In Byers and Pringle (2024), the ability of a single invasive species was quantified for a coastline with no alongshore variation in ocean currents. The population growth rate of the introduced species, when that species was at low abundance, was quantified as R, hereafter referred to as R^BP^ to avoid confusion. This R^BP^ subsumed all competition with native species, loss of larvae to regions outside of the habitat, and mortality before recruitment. If the invasive species is at low population, its population at generation 1, P^t=1^, would be related to its population at generation 0, P^t=0^, by

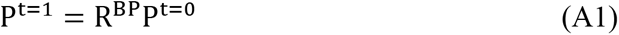

where P is the total population averaged over a large region.

In this work, we assume all species have the same dispersal behavior and mortality in the larvae, but that the native species release R_native_ larvae per adult and the introduced species releases R_introduced_ larvae per adult. The fraction of each species in the next generation will be proportional to the fraction of each species’ larvae which reach the habitat. Assuming for convenience that the total population is at the carrying capacity and that carrying capacity *P_max_* is 1.0, and that the population of the introduced species at time t=0 is *ε* and the native species is 1 − *ε*, the number of larvae *L* produced by each species is

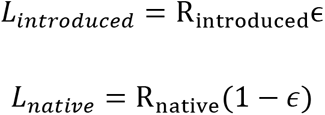

and the population of the introduced species in the next generation will be

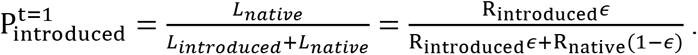

Expanding this equation with a Taylor series around *ε* = 0, which is equivalent to assuming that the population of the introduced species is much less than the carrying capacity, leads to

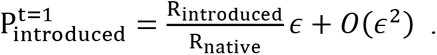

Comparing this with equation (A1) leads to the conclusion that

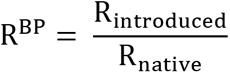

in the relevant limit of a small population of invaders relative to the carrying capacity.

In this manuscript, the relative competitiveness of the two species producing larvae with R’s R_native_ and R_introduced_ larvae per adult, and assuming the same dispersal and mortality in plankton, is defined with the parameter

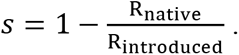

From this it can be shown that *s*=1-1/R^BP^ and R^BP^=1/(1-*s*).

Byers and Pringle (2024) argued that a novel introduced species could persist if

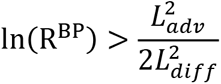

Where *L_adv_* is the mean downstream distance a larvae is transported, and *L_diff_* is the standard deviation of the transport distance of all the larvae. Using the results from above, this can be written in the notation of this paper as

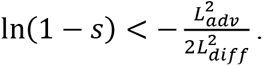

## Data availability statement

The analysis software used to make these results, and a guide to replicating the results presented here for different larval behaviors, phenologies, and regions, can be found at https://github.com/JamiePringle/PringleLushByers_InvasibilityInRealOcean and DOI:10.5281/zenodo.15814607. An interactive version of Figure **3** which includes the direction of mean larval transport can be found at https://jamiepringle.github.io/PringleLushByers_InvasibilityInRealOcean/.

